# Production of single-cell-protein (SCP) / poly(3-hydroxybutyrate-co-3-hydroxyvalerate) (PHBV) matrices through fermentation of archaea *Haloferax mediterranei*

**DOI:** 10.1101/2023.12.15.571855

**Authors:** Razan Unis, Rima Gnaim, Mrinal Kashyap, Olga Shamis, Nabeel Gnayem, Michael Gozin, Alexander Liberzon, Jallal Gnaim, Alexander Golberg

## Abstract

The idea of *in-situ* integrating poly(3-hydroxybutyrate-co-3-hydroxyvalerate) (PHBV) sieves in a single-cell protein (SCP) represents a promising approach to enhance the properties of microbial biomass as protein alternatives. Archaea SCP/PHBV matrix was successfully produced with a concentration of 8.0 ± 0.1 g L^-1^ and a productivity of 11.1 mg L^-1^ h^-1^ using *Haloferax mediterranei*. This was achieved by employing 30 g L^-1^ of enzymatically hydrolyzed bread waste (BW) and 200 g L^-1^ of red sea salt at 42 °C and with shaking at 150 rpm for 3 days. The amino acid profile of the SCP/PHBV matrix revealed a total amino acid content of 358 g kg^-1^, including 147 g kg^-1^ of essential amino acids. The protein quality of the *H. mediterranei* SCP/PHBV matrix was assessed using the *in-vitro* enzyme digestion method, indicating a high-quality protein with an *in-vitro* digestibility value of 0.91 and a protein digestibility-corrected amino acid score (PDCAAS) of 0.78. The PHBV component (36.0 ± 6.3% w/w) in the SCP/PHBV matrix consisted of a copolymer of 3-hydroxybutyrate and 3- hydroxyvalerate in a 91:9 mol% ratio, respectively. The simultaneous production of PHBV polymeric sieves within the *H. mediterranei* SCP/PHBV matrix provides an alternative protein source with enhanced physicochemical and thermal properties.

**Highlights:**

- SCP/PHBV matrices were produced from wasted bread by archaea *H. mediterranei*.
- This is the first report that explored the production and properties of SCP/PHBV.
- The presence of PHBV in SCP affected its physicochemical and thermal properties.
- SCP/PHBV with high-quality protein was achieved with a PDCAAS value of 0.78.

**Graphical abstract:** 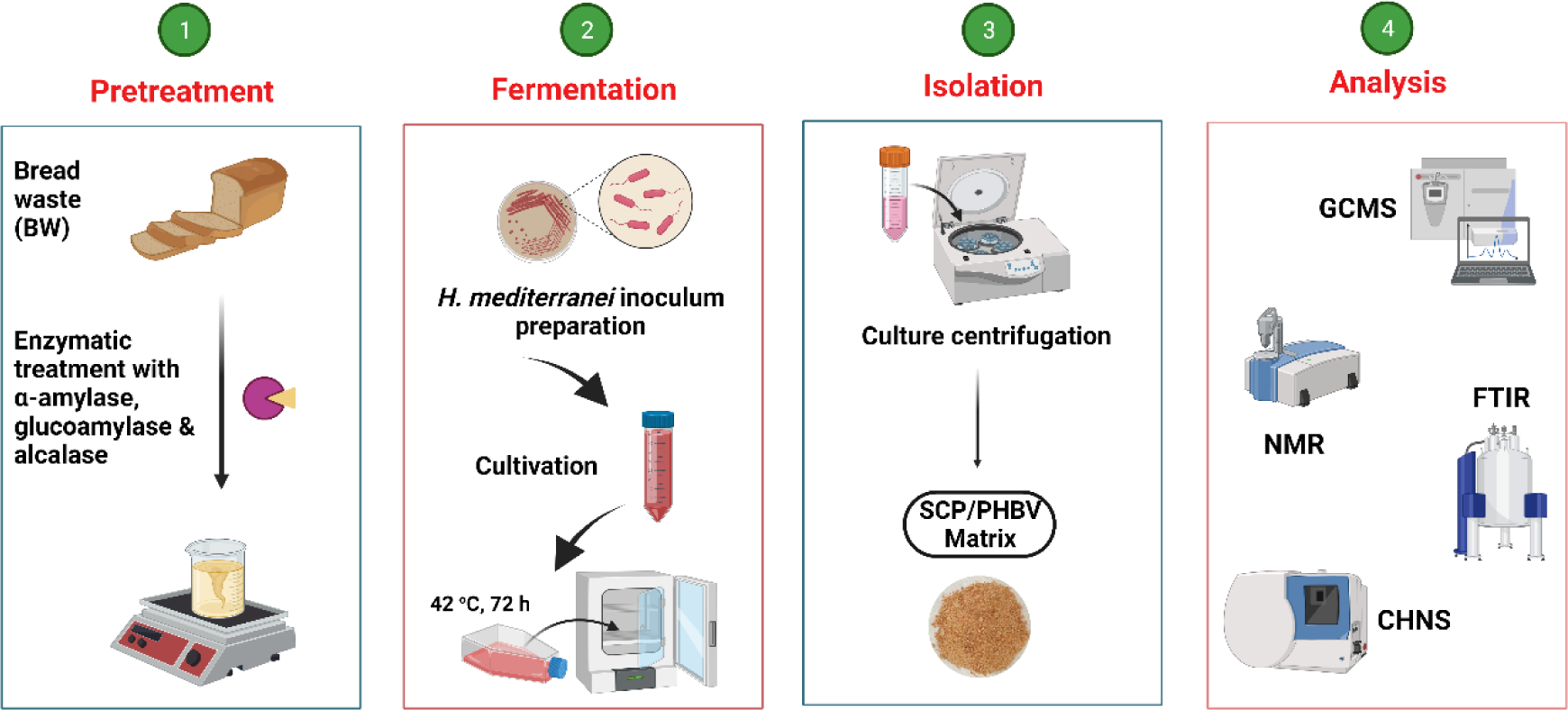

## 1. Introduction

The global population’s rapid growth and the consequential rise in starvation, malnutrition, and related diseases worldwide have intensified the need for adequate and continuous sources of nutrition (Bratosin, Darjan, & Vodnar, 2021). Meeting the growing demand for protein-rich foods through agriculture alone is challenging and complex (Fasolin et al., 2019); therefore, the search for alternative protein sources, such as microbial biomass, has been increasing (Al-Mudhafr, 2019).

Microorganisms have been employed for an extended period in producing food items with elevated protein content, including cheese and fermented soybean products (Upadhyaya, Tiwari, Arora, & Singh, 2016). The selection of microorganisms for this purpose depends on multiple criteria, such as rapid growth on a wide range of appropriate substrates (Sharif et al., 2021). Additional measures include nutritional (e.g., energy value, protein content and yield, amino acid, and essential amino acid balance) and procedural aspects (e.g., the sort of culture and the isolation method) (Ravindra, 2000). The main strategies regarding a substrate needed to produce microbial biomass containing high protein quality involve either utilizing low-grade waste materials or opting to use a simple carbohydrate source (Dunuweera, Nikagolla, & Ranganathan, 2021).

The protein harvested from microbial sources is known as “Single Cell Protein” (SCP) (Nasseri, Rasoul-Amini, Morowvat, & Ghasemi, 2011). SCP is the dried biomass of microorganisms that feed on various carbon sources (Jach, Serefko, Ziaja, & Kieliszek, 2022); it contains high protein levels in addition to fats, carbohydrates, vitamins, and minerals (Patelski et al., 2015; Khan, Khan, Ahmed, & Tanveer, 2010).

The production of SCP typically involves utilizing the following microbial genera (Salazar-López et al., 2022): (*i*) Bacteria are used extensively for the commercial scale production of SCP, with various strains including *Bacillus* (Zha et al., 2021) and *Pseudomonas* (Saejung, & Salasook, 2018). Bacteria have certain advantages over other microorganisms, owing to their high growth rate, short generation time (about 2 h), high protein content (80% of the total dry weight), and the abundance of sulfur amino acids, which require less synthesis energy (Onyeaka et al., 2022). Interestingly, nucleic acids, particularly RNA, constitute 15-16% of bacteria’s dry weight (Bratosin et al., 2021). (*ii*) *Saccharomyces* (Tropea et al., 2022) and *Candida* (Carranza-Méndez et al., 2022) are commonly used for SCP production using yeasts; they have been utilized for centuries in various industries, such as food and feed production, including baking, brewing, and winemaking (Ghorai et al., 2009). Yeasts possess the unique ability to grow in acidic conditions (pH 4.5-5.5); however, this is unsuitable for bacterial growth, thus eliminating the need for strict aseptic conditions (Dunuweera et al., 2021). Moreover, yeasts have several practical features, including the ability to use diverse substrates, genetic variation susceptibility, aggregation capacity, and high nutritional value (Johnson & Echavarri-Erasun, 2011). However, one disadvantage of using yeasts for SCP production is the limited availability of sulfur-containing amino acids in the obtained SCPs (Sharif et al., 2021). (*iii*) *Fusarium* (Aziz & Mohsen, 2002), *Aspergillus* (Azam, Khan, Ahmad, Khan, & Ali, 2014), and *Penicillium* (Valentino, Ganado, & Undanet, 2016) are commonly used filamentous fungi for SCP production. These fungi have a slow growth rate and are highly susceptible to yeast contamination, thus necessitating sterile growth conditions (Junaid, Khawaja, & Ali, 2020). (*iv*) *Spirulina* (Hu, 2004), *Chlorella* (Iwamoto, 2003), and *Scenedesmus* (Xia, Zhou, Zhang & Hu, 2013) are used for SCP production using algae. Algae, including *Cyanobacteria* and unicellular eukaryotes, are autotrophic and generate food by harnessing sunlight or artificial light, with CO2 as the carbon source (Sirohi, Lee, Yu, Roh, & Sim, 2021).

Algae have high digestibility (85-95%) due to their thin cell wall and low nucleic acid content, and they contain a high proportion of digestible proteins (Nalage et al., 2016). The archaea *Haloferax mediterranei* belongs to the extremely halophilic class of halobacteria and has numerous advantages for SCP production, such as the ability to survive and grow in high salinities, thus reducing microbial contamination risks (Pacholak, Gao, Gong, Kaczorek, & Cui, 2021). *H. mediterranei* has rapid growth compared with related organisms and a broader substrate spectrum (Mironescu, Ignatova, & Posten, 2004). It allows for adaptability to varying environmental conditions encountered during the fermentation process, encompassing changes in oxygen levels, temperature fluctuations, nutrient concentrations, and pH (Matarredona et al., 2021). In addition, *H. mediterranei* is sensitive to hypotonic media and can be efficiently lyzed in distilled water; therefore, using large quantities of organic solvents in the extraction process can be avoided (Keshavarz & Roy, 2010).

Halophiles have been investigated for functional protein production, specifically for enzymes such as pullulanases, proteases, lipases, hydrolases, amylases, and DNases (De Lourdes Moreno, Pérez, García, & Mellado, 2013), PHA-associated regulatory protein PhaR, granule-associated protein, PhaP (Zhang, Lin, & Chen, 2018), and extracellular polymeric substances (EPS) (Cui, Gong, Shi & Wang, 2017). In addition, Lillo et al. (Lillo & Rodriguez-Valera, 1990) have confirmed the suitability of halophiles as efficient protein bioreactors. In addition, *H. mediterranei* is a versatile intracellular polyhydroxyalkanoate (PHA) producer that has gained attention due to its potential to synthesize PHAs from simple and inexpensive carbon sources (Koller, 2015). PHAs are biopolymers that exhibit mechanical and thermal characteristics analogous to traditional plastics such as polyethylene and polypropylene. Among the interesting products produced by extremely halophilic archaea are poly(3- hydroxybutyrate) (PHB) and poly(3-hydroxybutyrate-co-3-hydroxyvalerate) (PHBV) (Gonzalez & Winterburn, 2022). PHB is characterized by its hardness and brittleness; it has a melting point closely approaching its degradation temperature, thereby limiting its utility due to a narrow temperature processing range (Ferre-Guell & Winterburn, 2018) However, PHBV is less crystalline, more flexible, and highly processable (Sato et al., 2021). Thus, it is gaining increasing importance in food packaging, agriculture, and biomedical applications such as tissue engineering scaffold fabrication, wound healing, and medical implant development (Cai et al., 2021). Interestingly, *H. mediterranei* is among the few microorganisms that synthesize PHBV from a simple and cheap carbon source without supplementing the 3-hydroxy valeric acid precursor (Zhao et al., 2013).

Numerous investigations on the production of PHA using *H. mediterranei* have focused on harnessing industrial and agricultural byproducts, such as extruded rice bran (Huang, Duan, Huang, & Chen, 2006), vinasse (Bhattacharyya et al., 2012), rice-based ethanol stillage (Bhattacharyya et al., 2015), cheese whey (Pais, Serafim, Freitas, & Reis, 2016), olive mill wastewater (Alsafadi & Al-Mashaqbeh, 2017), molasses wastewater (Cui, Gong, Shi & Wang, 2017), macroalgal biomass (Ghosh et al., 2019), ricotta cheese (Raho et al., 2020), date palm fruit waste (Alsafadi, Ibrahim, Alamry, Hussein, & Mansour, 2020), and bread waste (Montemurro et al., 2022). However, the industrial production of PHA is still hindered by the costly feed materials (Koller et al., 2005).

The co-production of PHA using various microorganisms (Li, Elhadi, & Chen, 2017) with other valuable chemicals has been demonstrated; these include amino acids (Gu et al., 2013), enzymes (Shamala, Vijayendra, & Joshi, 2012), alcohols (Xin, Zhang, Zhang, & Xia, 2007), molecular hydrogen (Singh, Kumar, Patel, & Kalia, 2013), biosurfactants (Rashid, Azemi, Amirul, Wahid, & Bhubalan, 2015), exopolysaccharides (Cui, Gong, Shi & Wang, 2017), and carotenoids (Kumar, Jun, & Kim, 2018). Umesh et al. investigated the production of PHA by *Bacillus subtilis* and SCP by *Saccharomyces cerevisiae* utilizing *Carica papaya* waste (Umesh, Priyanka, Thazeem, & Preethi, 2017). *Cupriavidus necator* cells were evaluated as a source of protein and used to recover PHA granules simultaneously (Chee, Lakshmanan, Jeepery, Hairudin, & Sudesh, 2019).

However, the coproduction of SCP/PHA matrix utilizing microorganisms, including archaea, has not been investigated regarding protein alternatives. The concept of integrating PHA-polymeric sieves into microbial SCP represents a promising avenue to enhance the properties of microbial protein alternatives. This combination could address a crucial aspect of sustainable food technology by leveraging the strengths of both components, i.e., protein and PHA, to overcome various limitations, such as texture, binding capacity, water- and oil-holding capacity, and elasticity, to produce protein substitutes (Alves et al., 2023).

The current research focused on assessing the potential of *H. mediterranei* as a versatile microorganism capable of simultaneously producing SCP/PHBV matrices. This was achieved by utilizing the enzymatic hydrolysate of bread waste (BW) as a nutrient-rich carbon/nitrogen source, along with red sea salt as a comprehensive growth medium. Specifically, the research aimed to (*i*) evaluate the variability in nutritional composition between different BW samples, including carbohydrates, lipids, and proteins, (*ii*) investigate the key factors affecting the growth of *H. mediterranei* under various conditions, including red sea salt and BW hydrolysate concentration, and pH variations, (*iii*) determine the composition of the resulting archaea SCP/PHBV matrices, including protein, PHBV polymer, ash content, macroelements, trace elements, lipids, and carbohydrates, (*iv*) determine the physical, chemical, thermal, and digestibility characteristics of the generated SCP/PHBV matrices, and (*v*) determine the stoichiometry of the fermentation process and its impact on reducing CH4 emissions and CO2 generation from BW.

## 2. Materials and methods

### 2.1. Materials

Yeast extract was obtained from Thermo Fisher Scientific (Difco™, Israel); it consisted (w/w) of 34.76% C, 9.30% N, 6.31% H, and 0.54% S with a C:N ratio of 3.74. Red sea salt was obtained from Aquazone Ltd. (Israel) and contained (g kg^-1^): Na 358.9, Cl 553.9, Mg 37.4, S 25.7, Ca 12.3, K 11.4, Sr 0.234, B 0.126, F 0.037, I 0.002, and other minor trace elements. The α-amylase enzyme from *Bacillus amyloliquefaciens* (≥ 250 U g^-1^), amyloglucosidase enzyme from *Aspergillus niger* (≥ 260 U mL^-1^), and alcalase® protease enzyme from *Bacillus licheniformis Subtilisin A* (≥ 2.972 U mL^-1^) were purchased from Sigma-Aldrich (Israel). Thirteen different BW samples and their mixtures were collected from local restaurants and bakeries (Kfar Qara’, Israel).

### 2.2. Dry weight, ash content, and elemental analysis of BW

#### 2.2.1. Dry weight of BW

Ten g of fresh BW samples (in triplicate) were cut into 1-2 cm pieces and were dried in an air oven (Carbolite, Israel) at 105 °C for 3 days. The dried samples were ground to a fine powder using a blender (Gold line, Israel), then weighed and stored in closed containers at −20 °C until use.

#### 2.2.2. Ash content of BW

One g of dry powder of BW (in triplicate) was put in a pre-weighed crucible. The crucible containing the BW sample was subjected to heating at 550 °C for 5 h. Next, the crucible with the remaining ash was cooled at 25 °C and weighed, and the ash content was calculated (Gnaim et al., 2023).

#### 2.2.3. Elemental analysis of BW

CHNS elemental analysis of the dry BW samples (in triplicate) was determined utilizing a Thermo Scientific™ FLASH 2000 CHNS/O Analyzer (Technion, Israel). The determination of both the macroelements and trace elements involved the utilization of a PerkinElmer NexION 2000 inductively coupled plasma mass spectrometer (ICP-MS). This analysis was carried out at the Field Service Lab Central District (Hadera, Israel).

### 2.3. Enzymatic hydrolysis of BW

Enzymatic hydrolysis of dry BW samples was carried out in 3 steps (Hudečková Šupinová, & Babák, 2017). First, 10 g of BW samples (in triplicate) were homogenized with 100 mL of distilled water. The pH of the slurry was adjusted to 6.0; then, 4 mL (250 U g^-1^) of thermostable α-amylase was added. The mixture was kept at 80 °C for 3 h with magnetic stirring at 150 rpm. The liquefaction was curtailed by freezing the mixture at −20 °C for 12 h. In the second hydrolysis step, saccharification, the pH of the liquefied suspension was adjusted to 4.2, and the saccharification was performed in liquefied suspension by adding 4 mL (260 U mL^-1^) amyloglucoamylase at 60 °C for 90 min. Heating the enzyme at 80 °C for 5 min resulted in its inactivation, followed by subsequent cooling of the mixture to room temperature. Next, 1 mL (2.972 U mL^-1^) of endopeptidase alcalase was used in the third step after pH adjustment to 8.0, followed by heating at 80 °C for 24 h. Then, the enzyme’s activity was nullified by subjecting the suspension to heating at 100 °C for 5 min. Finally, the mixture was filtered and stored at 4 °C until use.

### 2.4. Archaea strain and red sea salt medium preparation

*H. mediterranei* (ATCC 33500, CCM 3361) from the DSMZ (DSM 1411) culture collection was used for strain activation and culture. The following medium was employed for all *H. Mediterranei* cultivation experiments under different conditions: 90-200 g L^-1^ of red sea salt powder, 0-24 mL L^-1^ of trace element solution, 0-0.7 g L^-1^ of NH4Cl, 0-0.6 g L^-1^ of KH2PO4, 0-55 g L^-1^ of glucose, 0-5 g L^-1^ of yeast extract, and 0- 55 g L^-1^ of BW hydrolysate were added to 800 mL deionized water with a stirring rate of 300 rpm at 42 °C for 3 h. Next, the solution was microfiltered under vacuum (Corning® 500 mL, USA), and deionized water was added to complete the volume up to 1000 mL. Finally, the pH of the solution was adjusted in the range of 2 to 13 and kept at 4 °C until use.

### 2.5. Cultivation of H. mediterranei in 96-well plates

Cell pellets obtained from 2-24 µL of inoculum solution were resuspended in 198- 176 µL of red sea salt medium in a 96-well plate, sealed with an adhesive plate sealer, and cultivated at 42 °C for 120 h with shaking at 150 rpm. The culture microplate was shaken for 30 seconds and then placed into the multi-reader (Infinite M Plex Elisa, Tecan, Austria). The optical density was measured at 600 nm at 25 °C at specific times of 0, 24, 48, 72, 96, and 120 h (5 replicates) with a cell-free supernatant serving as a blank (in triplicate). After 120 h, the cultivation solutions were transferred to 2 mL microcentrifuge tubes (Tarasons, India), and the cell biomass was collected by centrifugation (Neofuge 13R high-speed refrigerated benchtop centrifuge, China), operating at 13000 rpm for 10 min. Next, the biomass was washed with 200 µL deionized water and dried at 60 °C for 24 h. The resulting biomass was weighed and analyzed for PHBV and protein content using FTIR.

### 2.6. Batch cultivation of H. mediterranei in BW and red sea salt medium

Batch cultivation (in triplicate) was conducted in culture flasks using a 100 mL solution containing 20 mL of *H. mediterranei* inoculum, 3 g of BW hydrolysate, and 20 g of red sea salt. The pH of the mixture was adjusted to 7.3 and then cultured at 42 °C for 72 h with constant shaking at 150 rpm. Following the cultivation, the cultures were subjected to centrifugation, washing, and drying at 60 °C for 24 h. Finally, the dried biomass was weighed, and the components, including protein and PHBV, were examined using Fourier transform infrared (FTIR).

### 2.7. PHBV isolation from the SCP/PHBV matrix

A mixture of 200 mg of *H. mediterranei* SCP/PHBV (in triplicate) and a 10 mL chloroform was subjected to reflux at 62 °C for 12 h. Next, the cooled mixture was filtered using a Whatman filter (no. 4, Macherey-Nagel, Germany). The off-white solid (non-PHBV cell mass) was collected, dried at 60 °C for 12 h and weighed. In parallel, the supernatant, i.e., a PHBV/chloroform solution, was gradually poured into 20 mL methanol. Finally, the PHBV precipitate was isolated by centrifugation at 4000xg for 30 min, dried at 60 °C for 24 h, weighed, and stored at −20 °C.

### 2.8. Analysis of amino acids in H. mediterranei SCP/PHBV matrix

First, 100 mg of *H. mediterranei* SCP/PHBV matrix (in triplicate) was placed in a glass tube and hydrolyzed with 5 mL of 6N HCl solution and phenol at 110 °C for 22 h. Another 100 mg aliquot of this sample (in triplicate) was first oxidized with formic acid and hydrogen peroxide at 2-8 °C for 16 h. Next, the oxidized samples were dried under vacuum and then hydrolyzed with 5 mL of 6N HCl and phenol at 110 °C for 22 h. Aliquots of the two hydrolysates were dried by a vacuum centrifuge and dissolved in an amino acid sample buffer. The hydrolysate solutions were sonicated, vortexed, and filtered using a 0.45 µm nylon filter. Next, 20 µl of the hydrolysate solutions were injected into the Biochrom 30+ Amino-Acid-Analyzer (AminoLab, Analytical Laboratory Services, Israel). The amino acids were separated on an ion exchange column (Biochrom H-1552), derivatized with ninhydrin after eluting from the column, detected at 570 and 440 nm, and quantified against a standard.

### 2.9. Density, ash content, and water- and oil-holding capacities of SCP/PHBV matrix

#### 2.9.1. Bulk density measurement of SCP/PHBV matrix

Seven g of SCP/PHBV powder (in triplicate) was placed in a 10 mL measuring cylinder and taped until no apparent volume reduction was observed. Next, the final volume of the SCP/PHBV powder was recorded. Finally, the bulk density was calculated and reported in units of g cm^−3^ (Razzaq, Khan, Maan, & Rahman, 2020).

#### 2.9.2. True density measurement of SCP/PHBV matrix

Seven g of SCP/PHBV powder (in triplicate) was placed in a 25 mL glass measuring cylinder containing 10.0 mL hexane. Next, the final volume of the mixture was recorded. The true density was calculated and expressed as g cm^−3^ (Razzaq et al., 2020).

#### 2.9.3. Water- and oil-holding capacities of SCP/PHBV matrix

First, 10 mL of distilled water or 10 mL of sunflower oil was added to 2.00 g of SCP/PHBV powder (in triplicate) in a 15 mL polypropylene centrifuge tube, then vortexed, and allowed to remain in a static position at 25 °C for 30 min. Finally, the suspension was centrifuged, and the tube containing the sediment was weighed (Razzaq et al., 2020).

### 2.10. Determination of the animal-safe accurate protein quality score

SCP/PHBV’s protein quality was assessed by evaluating their amino acid composition and *in-vitro* protein digestibility-corrected amino acid score (PDCAAS) using the Megazyme assay kit (Wicklow, Ireland, https://www.megazyme.com/). To 500mg of milled SCP/PHBV sample (in triplicate), 19 mL of 0.06N HCl was added, and the mixture was incubated at 37 °C for 30 min with shaking at 120 rpm. Next, 1 mL of pepsin solution was added, and the sample was incubated at 37 °C for 1 h. The pH was brought to 7.4 with 2 mL of 1M Tris buffer, followed by the addition of 200 μL of a trypsin-chymotrypsin mixture. The sample was vortexed and incubated at 37 °C for 4 h with shaking at 120 rpm, then placed in a boiling water bath for 10 min. Next, the sample was vortexed, cooled to 25 °C for 20 min, mixed with 1 mL of 40% trichloroacetic acid solution, incubated at 4 °C overnight, and then centrifuged at 25 °C for 10 min at 13,800 rpm. A 10-fold dilution in acetate buffer (50 mM, pH 5.5) was performed before the colorimetric assay. PDCAAS values were computed by utilizing the Megazyme Mega-CalcTM programme (K-PDCAAS Mega-Calc). A standard curve derived from L-glycine, used to plot the absorbance values recorded at 570 nm against L-glycine concentrations spanning from 0 to 1 mM, was used to assess the primary amine concentration (*CI*) in unidentified samples. The concentration of primary amines (in mM) in the unknown samples was determined using equation (1), in which *CI* represents the unknown primary amine concentration, *Y* corresponds to the absorbance, *B* denotes the *y*-intercept, and *A* represents the slope of the line.

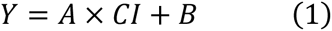

Equation (2) was used to determine the primary amine concentration (*C2*) in the initial sample solution. In this equation, *CI* represents the concentration of primary amines in the samples after dilution, *D* stands for the dilution factor applied to the samples before amine measurement, 1.25 denotes the dilution factor associated with trichloroacetic acid, *W* represents the weight of the sample (g), and 0.5 signifies the nominal quantity (g).

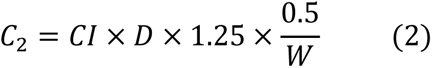

Amino acid constants were employed to compute the adjusted primary amine concentration (*CN*) for the individual amino acids, as indicated in equation (3). In this equation, *C2* represents the adjusted primary amine concentration in the initial sample solution (measured in mM), whereas proline, lysine, histidine, and arginine denote the concentrations of these respective amino acids in the original sample. The constants 2, 0.5, 0.2 and 0.2 are specific values associated with the corresponding amino acids.

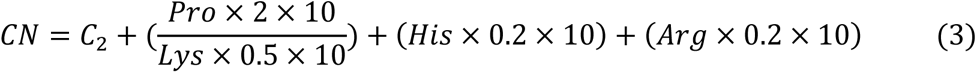

*In-vitro* digestibility was determined utilizing equation (4), which relies on established literature values for the rat model. To assess the corrected primary amine concentration (*CN*), the data fits were compared using a linear regression equation. In this equation, *X* denotes the corrected primary amine concentration for individual samples, *M* represents the slope of the regression line, *B* signifies the *y*-intercept, and 100 is the conversion factor used to convert percentages to grams.

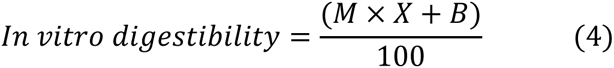

The amino acid ratio and the identification of the limiting amino acids, expressed in grams per 100 grams of protein, were computed using the total crude protein values in the dry SCP/PHBV, as demonstrated in equation (5). The amino acid ratio within the sample was determined in accordance with the recommended values, as outlined in equation (6). The *in-vitro* PDCAAS score was derived by multiplying the *in-vitro* digestibility obtained from equation (4) by the limiting amino acid ratio (the smallest value) obtained from equation (6).

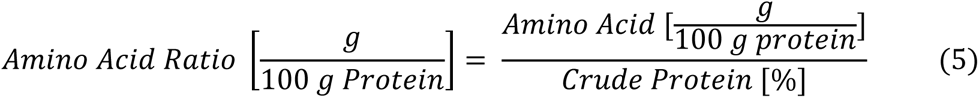

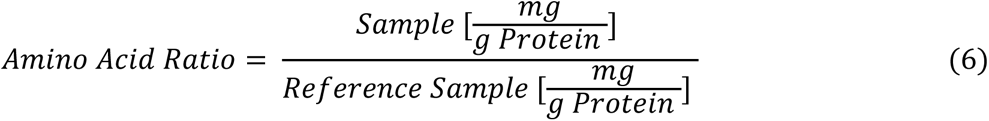

### 2.11. Analysis methods

#### 2.11.1. Thermogravimetric and differential scanning calorimetry analysis

Thermogravimetric analysis (TGA) was employed to determine SCP/PHBV’s degradation temperature (*Td*), along with the extracted PHBV and protein constituents. This involved subjecting the samples to a controlled heating regimen spanning the temperature range from 30 to 600 °C, with a constant heating rate of 10 °C min^-1^. Furthermore, the melting temperature (*Tm*) was ascertained utilizing differential scanning calorimetry (DSC) over a temperature range of 30-600 °C, by employing a heating rate of 10 °C min^-1^ (NETZSCH STA 449F5 STA449F5A-0214-M).

#### 2.11.2. Fourier transform infrared analysis

FTIR spectra of the SCP/PHBV matrix, isolated PHBV, and isolated protein were recorded by employing a Thermo Scientific™ Nicolet™ iS50 FTIR Spectrometer, covering a spectral range from 400 to 4000 cm^−1^ through 16 scan repetitions.

#### 2.11.3. Nuclear magnetic resonance analysis of PHBV

The nuclear magnetic resonance (^1^H- and ^13^C-NMR) spectra were acquired on a Bruker 500 MHz NMR spectrometer while dissolving 15 mg of isolated PHBV in 0.5 mL of CDCl3 while heating at 50 °C for 10 min.

#### 2.11.4. Gel permeation chromatography analysis of PHBV

The gel permeation chromatography (GPC) analysis was carried out using an Agilent 1260 Infinity II GPC system equipped with security Guard Cartridges GPC 4 x 3.0 mm ID 3/Pk XAJ0-9292, two GPC LF-804 columns, and one KF-803 column arranged in a sequence. The analysis was conducted in tetrahydrofuran at 35 °C, following the methodology recently reported (Gnaim et al., 2023).

#### 2.11.5. Gas chromatography-mass spectrometry

The characterization and quantification of the isolated PHBV were performed via gas chromatography-mass spectrometry (GC-MS) after the direct acid-catalyzed methanolysis of SCP/PHBV, as described recently (Gnaim et al., 2021).

### 2.12. Statistical analysis

Results were evaluated using a one-way ANOVA for the following variables: dry weight, ash content, and the C:N ratio of BW. Additionally, a two-way ANOVA with repeated measures was performed to investigate several abiotic factors (the pH, BW concentration, BW type, and red sea salt concentration) that affected the growth of *H. mediterranei*. Statistically significant differences were identified using Tukey multiple comparison tests using GraphPad Prism version 10, with significance, indicated when *p <* 0.05.

## 3. Results and discussion

### 3.1. Collection and analysis of BW samples

Various BW samples (BW-1 to BW-13) were collected from local restaurants and bakeries (Supplementary Figure 1S). Their commercial nutritional values, including carbohydrates, lipids, and proteins, are listed in Table 1. The carbohydrate contents in the BW samples ranged from 4 to 56.5% (w/w), lipids from 0 to 25.4% (w/w), and proteins from 9.1 to 27.5% (w/w). In addition, all BW samples were analyzed for their dry weight, ash content, C:N ratio, and the N-to-protein conversion factor. The corresponding values for dry weight ranged from 53.8 to 65.9% (w/w), ash content from 1.6 to 7.1% (w/w), C:N ratio from 7.1 to 23.4, and N-to-protein conversion factor from 4.9 to 7.9. The average N-to-protein conversion factor obtained for all BW samples in this study was 6.1, similar to that reported for wheat (5.8) (Fujihara, Sasaki, Aoyagi, & Sugahara, 2008). The protein concentration of BW was estimated by multiplying its nitrogen content, which was determined by elemental analysis, by the average N-to-protein conversion factor (6.1). Among the BW samples analyzed by one-way ANOVA and Tukey’s multiple comparisons test, BW-9 displayed the highest dry weight (65.6 ± 0.6% w/w) (significance, *p* < 0.0001 with BW-1 to BW-4 and BW-7 to BW-13 but not significant with BW-5, *p* = 0.4103, and BW-6, *p* = 0.1092). BW-2 exhibited the highest ash content (7.1 ± 0.4% w/w) (significance, *p* < 0.0001 with BW-1 to BW-7 and BW-9 to BW-13, but not significant with BW-8, *p* = 0.2287). BW-9 exhibited the highest C:N ratio (23.4 ± 0.1 w/w) (significance, *p* < 0.0001 with all BW samples). BW-4 presented the highest N-to-protein conversion factor (7.9 ± 0.1). The large variability between different BW samples regarding the values of carbohydrates, lipids, proteins, dry weight, ash content, and the C:N ratio could result from differences in moisture content, mineral composition, protein composition, and nitrogen content in the BW samples. However, all BW samples were found to be rich in nitrogen, i.e., having a C:N ratio from 7.1 to 23.4, which is an essential factor for the growth of *H. mediterranei* and is suitable for microbial protein production.

**Table 1.** The carbohydrates (% w/w), lipids (% w/w), proteins (% w/w), dry weight (DW, % w/w of fresh weight), ash content (AC, % w/w of DW), CNS elements (% w/w of DW), C:N ratio, and N to protein conversion factor in bread waste (BW) samples. The BW samples were dried at 105 °C for 3 days. Mean ± SD, *n* = 3.

| BW sample | BW type | Commercial nutritional values |  |  | DW, % w/w<br>105 °C, 3 days | AC<br>% w/w<br>of DW | Elemental analysis (in DW) |  |  | N-to-protein<br>conversion<br>factor |
| --- | --- | --- | --- | --- | --- | --- | --- | --- | --- | --- |
|  |  | Carbohydrates<br>% w/w | Lipids<br>% w/w | Proteins<br>% w/w |  |  | C<br>% w/w | N<br>% w/w | C:N<br>ratio |  |
| BW-1 | Whole wheat | 36.2 | 2.8 | 10.3 | 58.8 ± 0.6 | 3.3 ± 0.1 | 43.1 ± 0.5 | 3.0 ± 0.1 | 14.4 ± 0.2 | 5.8 ± 0.1 |
| BW-2 | Sliced wholemeal bread | 25.0 | 3.1 | 15.1 | 53.8 ± 0.5 | 7.1 ± 0.4 | 43.8 ± 1.1 | 4.5 ± 0.0 | 9.7 ± 0.2 | 6.2 ± 0.0 |
| BW-3 | Light bread | 25.5 | 0.9 | 13.0 | 62.8 ± 0.6 | 4.6 ± 0.4 | 39.9 ± 4.1 | 3.8 ± 0.3 | 10.5 ± 0.1 | 5.5 ± 0.3 |
| BW-4 | Sliced black bread | 56.5 | 2.5 | 10.3 | 61.3 ± 0.8 | 2.9 ± 0.3 | 41.8 ± 0.1 | 2.1 ± 0.0 | 19.9 ± 0.1 | 7.9 ± 0.1 |
| BW-5 | Light bite pita | 31.5 | 0.0 | 10.2 | 64.5 ± 0.7 | 4.0 ± 0.0 | 43.6 ± 0.4 | 3.2 ± 0.0 | 13.6 ± 0.1 | 4.9 ± 0.1 |
| BW-6 | Flax bread | 4.0 | 16.4 | 27.5 | 64.1 ± 0.4 | 3.7 ± 0.3 | 52.0 ± 1.6 | 7.3 ± 0.2 | 7.1 ± 0.2 | 5.9 ± 0.2 |
| BW-7 | Sliced uniform white | 51.2 | 1.9 | 9.5 | 61.0 ± 0.7 | 1.6 ± 0.1 | 42.6 ± 0.9 | 2.4 ± 0.1 | 17.8 ± 0.3 | 6.5 ± 0.1 |
| BW-8 | Cereal bread | 25.4 | 25.4 | 12.2 | 59.7 ± 0.4 | 5.7 ± 1.2 | 42.7 ± 0.9 | 3.0 ± 0.2 | 14.2 ± 0.3 | 6.8 ± 0.2 |
| BW-9 | White pita-1 | 55.7 | 1.2 | 9.1 | 65.6 ± 0.6 | 3.4 ± 0.6 | 42.0 ± 1.3 | 2.1 ± 0.1 | 20.0 ± 0.6 | 6.6 ± 0.1 |
| BW-10 | Rye flour pita | 46.0 | 2.0 | 11.2 | 54.5 ± 0.5 | 3.2 ± 0.1 | 42.4 ± 0.2 | 3.0 ± 0.2 | 14.1 ± 0.1 | 6.9 ± 0.2 |
| BW-11 | Wholemeal pita | 58.0 | 2.0 | 10.0 | 58.1 ± 0.4 | 3.5 ± 0.2 | 42.2 ± 0.3 | 3.0 ± 0.0 | 14.1 ± 0.1 | 5.7 ± 0.1 |
| BW-12 | Light pita | 43.0 | 0.7 | 9.7 | 56.9 ± 0.3 | 4.4 ± 1.4 | 42.9 ± 0.8 | 3.2 ± 0.1 | 13.4 ± 0.3 | 5.3 ± 0.1 |
| BW-13 | White Pita-2 | 55.7 | 1.2 | 5.8 | 58.8 ± 0.4 | 3.3 ± 0.6 | 39.7 ± 0.2 | 1.7 ± 0.0 | 23.4 ± 0.1 | 5.8 ± 0.1 |

### 3.2. Optimization of SCP/PHBV matrix production from BW by H. mediterranei

The current study extensively investigated the key factors that impact the growth of *H. mediterranei* under various challenging conditions (Fig. 1). Specifically, the effect of the concentration of red sea salt (90-200 g L^-1^), trace elements (0-24 g L^-1^), NH4Cl (0-7 g L^-1^), KH2PO4 (0-6.6 g L^-1^), glucose (0-55 g L^-1^), and BW hydrolysate (5-55 g L^-1^), as well as pH variations (2-13) were investigated. The enzymatic hydrolysis of BW samples and their mixtures was carried out with liquefaction by α-amylase, saccharification by amyloglucoamylase, and protein hydrolysis by endopeptidase alcalase. *H. mediterranei* cultivation was carried out in a 96-well plate at 42 °C and at 150 rpm for 120 h. The outcome of these experiments indicated that the pH, red sea salt, and BW concentrations had the most significant effects among all the parameters studied.

**Fig. 1.**
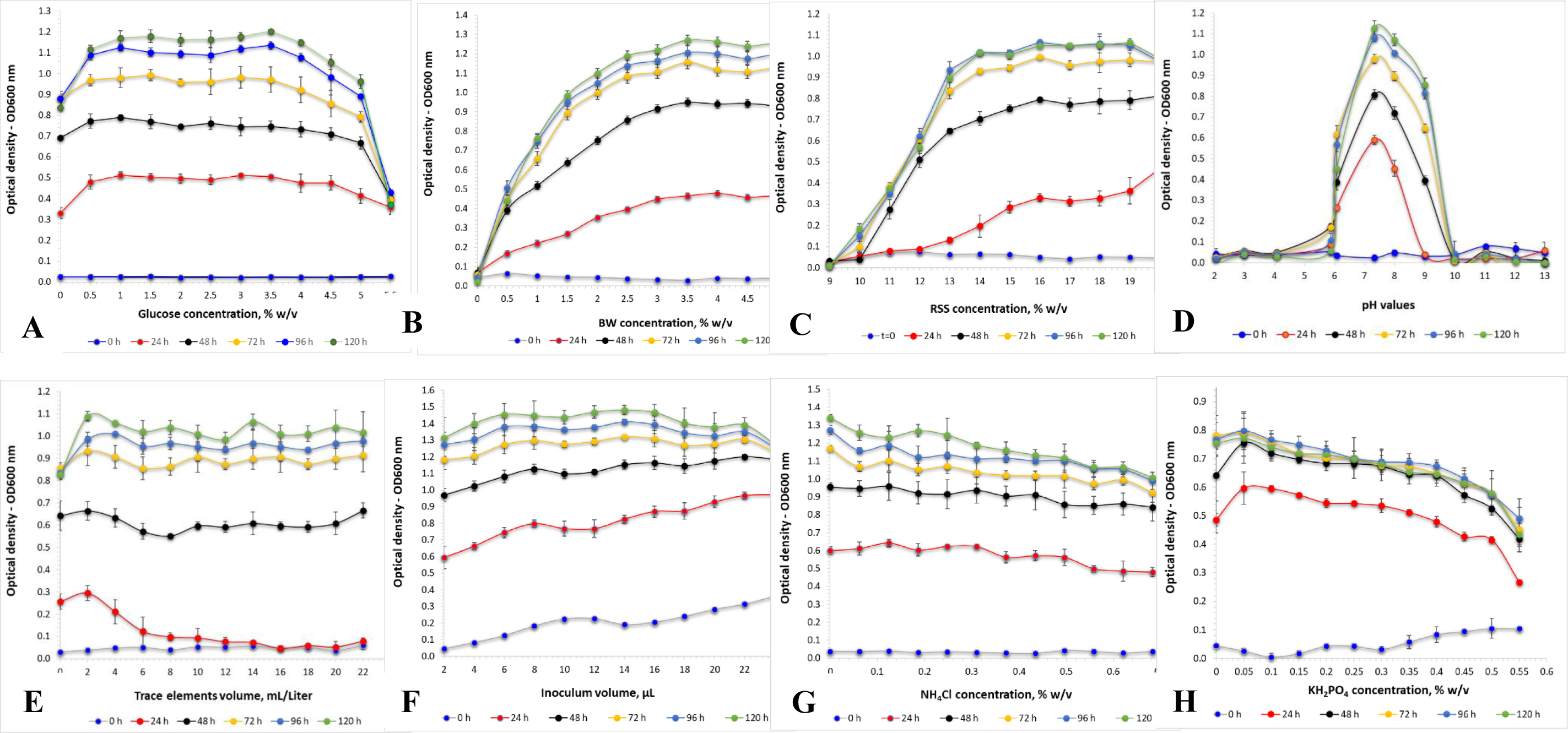
(A) Effect of different glucose concentrations on the optical density (OD600) of *H. mediterranei* culture. *H. mediterranei* was cultivated using a 96-well plate, 198 µL medium, 2 µl inoculum, 5 replicates for each treatment, 3 triplicates for each blank, pH 7.5, yeast extract 0.5% w/v, red sea salt 17.5% w/v, 150 rpm, at 42 °C. (B) Effect of different concentrations of bread waste (BW). (C) Effect of different concentrations of red sea salt. (D) Effect of different pH values. (E) Effect of different trace element concentrations. (F) Effect of different inoculum volumes. (G) Effect of different NH4Cl concentrations. (H) Effect of different KH2PO4 concentrations.

#### 3.2.1. Effect of BW concentration on the SCP/PHBV production

A statistically significant difference (*p* < 0.0001) in *H. mediterranei* growth was detected between the BW concentration ranges of 5 to 20 g L^-1^ and 25 to 55 g L^-1^ (Fig. 1B). Optimized archaea growth was observed using a concentration range of 25 to 55 g L^-1^ without statistical significance (*p* > 0.05). The optimized specific growth rate (µ) and doubling time (d.t.) (Rubin, Gnaim, Levi, & Zucker 2023) obtained with 30 g L^-1^ BW were 0.11 ± 0.02 h^-1^ and 6.3 ± 0.9 h, respectively. The composition of the resulting archaea SCP/PHBV matrix was determined by FTIR analysis, which revealed a PHBV content of 21.4 ± 1.1 % (w/w) and a protein content of 24.4 ± 1.2 % (w/w). Montemurro et al. previously demonstrated the suitability of enzymatically hydrolyzed BW, supplemented with seawater, as a substrate for bioplastic production by fermenting *H. mediterranei* (Montemurro et al., 2022).

#### 3.2.2. Effect of red sea salt concentration on SCP/PHBV production

Different concentrations of red sea salt (Fig. 1C) were examined in the cultivation of *H. mediterranei*. No statistically significant differences in the OD600 (*p* > 0.05) were observed for red sea salt concentrations in the range of 150 to 200 g L^-1^. On the other hand, at 100 g L^-1^, there was no observed growth, and a delayed growth response was detected at lower concentrations of red sea salt, with significant differences (*p* < 0.05) between the concentration ranges of 90 to 140 g L^-1^ and 150 to 200 g L^-1^. The optimized values of 0.097 ± 0.002 h^-1^ for µ and 7.2 ± 0.2 h for d.t. were achieved using 200 g L^-1^ red sea salt. The SCP/PHBV matrix produced under these conditions contained 20.0 ± 1.0% (w/w) of PHBV and 20.5 ± 1.0 % (w/w) of protein. These findings align with the Matarredona et al. study, which reported an optimum growth rate at sea salt concentrations between 100 and 325 g L^-1^ (Matarredona et al., 2021). *H. mediterranei* requires a minimum of 100 g L^-1^ salt for growth and can thrive in its natural environment with salt concentrations above 200 g L^-1^. This indicates the remarkable ability of *H. mediterranei* to withstand extreme salinity levels, and it has efficient osmoregulatory mechanisms that most likely allow it to maintain an adequate cellular water balance and grow effectively under varying salinity conditions.

#### 3.2.3. Effect of pH on SCP/PHBV production

A wide range of medium pH values, from 2 to 13, was applied in cultivating *H. mediterranei*. It was observed that *H. mediterranei* could not survive when the pH of the cultivation medium was lower than 5.9 or higher than 9. However, it exhibited its most significant growth (*p* < 0.0001) at a pH of 7.3 with a µ value of 0.133 ± 0.005 h^-1^, a d.t. value of 5.2 ± 0.2 h, a PHBV value of 19.4 ± 1.0 % w/w, and a protein concentration of 21.7 ± 1.1 % w/w (Fig. 1D). The results were similar to those of Matarredona et al., where the optimum growth of *H. mediterranei* was achieved at pH 7.25 (Matarredona et al., 2021).

#### 3.2.4. Effect of the BW type on the SCP/PHBV production

A two-way ANOVA study with repeated measures was performed to examine the effect of BW type on *H. mediterranei* growth (Fig. 2). Following this analysis, a Tukey multiple comparison test was conducted to pinpoint significant differences between the group means; it revealed a significant effect of time, e.g., 24 h compared to 48 h, F (1, 84) = 6819, *p* < 0.0001. Moreover, it was observed that there was no significant effect of the various bread samples on the growth of *H. mediterranei*, as indicated by the non-significant results of the statistical analysis (F (11, 84) = 1.147, *p* = 0.3362). Additionally, the interaction between the BW sample type and time was significant, F (11, 84) = 6.234, *p* < 0.0001. Subsequently, a Tukey multiple comparisons test was conducted to investigate pairwise differences between the BW sample levels. The results revealed no significance (*p* > 0.05) between all BW samples.

**Fig. 2.**
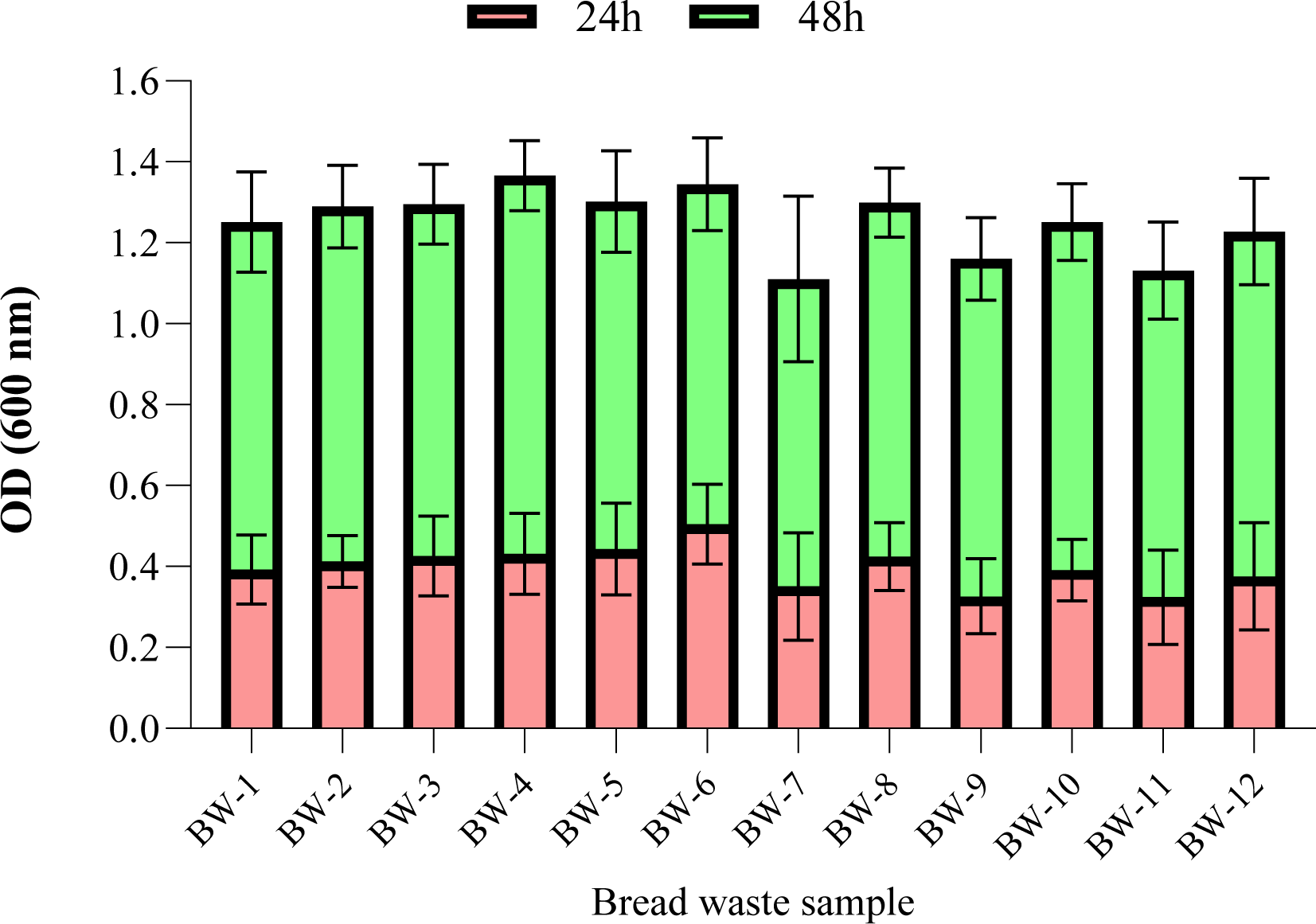
Effect of different bread waste samples (BW-1 to BW-12) (20 g L^-1^) on the optical density (OD600) of *H. mediterranei* culture. *H. mediterranei* was cultivated using a 96-well plate, 198 µL medium, 2 µl inoculum, 5 replicates for each treatment, triplicate for each blank, pH 7.5, red sea salt 17.5% w/v, 150 rpm, at 42 °C, for 24 h (red) and 48 h (green).

In summary, the study highlighted the significance of substrate concentration, specifically the enzymatic hydrolysate of BW, with an optimum performance observed at 30 g L^-1^ of BW. Salinity, represented by red sea salt concentrations of 150 to 200 g L^-1^, substantially influenced the growth and provided the most favorable conditions. Moreover, pH emerged as a vital factor, and neutral values at around 7.3 resulted in the highest growth rates. Furthermore, the study’s findings enable the use of different BW combinations, effectively minimizing the impact of BW composition fluctuations on the quality of the resulting archaea SCP/PHBV.

### 3.3. Batch cultivation of H. mediterranei for SCP/PHBV production

In the batch cultivation of *H. mediterranei* under optimized conditions, i.e., pH 7.3, at 42 °C, and 150 rpm for 3 days, a mixture of 30 g L^-1^ of BW hydrolysate and 200 g L^-1^ of red sea salt was found to produce a maximum SCP/PHBV concentration of 8.0 ± 0.1 g L^-1^ with a productivity of 11.1 mg L^-1^ h^-1^. The SCP/PHBV matrix contained 36.0 ± 6.3% (w/w) of PHBV. The stoichiometry for the fermentation of *H. mediterranei* on BW was obtained by a CHNS elemental mass balance, revealing a chemical formula of [C355H660N23S2.6O297] for BW and [C241H408N27S3O390] for the SCP/PHBV matrix, as presented in equation (7). According to the fermentation reaction, 649 g of CO2 can be generated from 1000 g of BW.

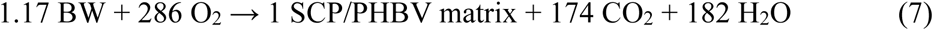

Examples of cell biomass and PHA production by *H. mediterranei* utilizing various industrial and agricultural wastes are presented in Table 2. These results show cell biomass values ranging from 0.46 to 65% (w/w). However, it is worth noting that in these previous studies, the PHA polymer was typically separated. In contrast, our study takes a novel approach by maintaining the PHA within the cell biomass, forming a SCP/PHBV matrix. This innovative approach could enhance the chemical, physical, and thermal properties of the SCP/PHBV matrices, thus providing an alternative protein source with improved characteristics.

**Table 2.** Literature examples of industrial and agricultural waste bioconversion to cell biomass and PHA by *H. mediterranei*.

| Waste | Cell biomass<br>g L<sup>-1</sup> | PHA<br>(% w/w) | Reference |
| --- | --- | --- | --- |
| Bread waste | 3-6 | 24 | Montemurro et al., 2022 |
| Rice bran | 63-140 | 27-56 | Huang et al., 2006 |
| Cheese whey | 5.7 | 29-65 | Pais et al., 2016 |
| Olive mill wastewater | 0.46 | 43 | Alsafadi & Al Mashaqbeh, 2017 |
| Macroalgal biomass | 3.8 | 57 | Ghosh et al., 2019 |
| Ricotta cheese | 10-18 | 6-10 | Raho et al., 2020 |
| Date palm fruit waste | 12.8 | 24 | Alsafadi et al., 2020 |

### 3.4. Composition of the archaea SCP/PHBV matrix

#### 3.4.1. Ash content

The ash content in the *H. mediterranei* SCP/PHBV matrix was 20.15 ± 1.94% w/w, which surpasses that of several protein sources, including yeast (5-9.5% w/w), bacteria (3-7% w/w), plants (0.5-1.6% w/w), milk (5.8% w/w), beef (4% w/w), and eggs (3.9% w/w) (Nalage et al., 2016), but it was lower than that of seaweed (30-40% w/w) (Sonchaeng et al., 2023). This notable mineral content of *H. mediterranei* is due to its high salinity habitat.

#### 3.4.2. CHNS elemental analysis

The *CHNS* elemental analysis of the SCP/PHBV matrix revealed 28.9 ± 4.9% carbon, 3.8 ± 0.6% nitrogen, and 0.9 ± 0.2% sulfur. The C:N ratio was 7.6 ± 1.3, and the N content determined by the Kjeldahl method was 4.0 ± 0.3%, implying a protein-rich composition.

#### 3.4.3. Macroelements and trace elements

The total macroelements in SCP/PHBV were 10.91 ± 0.13 g kg^-1^ and mainly consisted of Na 7.05 ± 0.36 g kg^-1^, K 1.04 ± 0.18 g kg^-1^, P 0.91 ± 0.04 g kg^-1^, Mg 0.85 ± 0.05 g kg^-1^, S 0.73 ± 0.10 g kg^-1^, and Ca 0.34 ± 0.02 g kg^-1^. The total trace elements in SCP/PHBV were 91.5 ± 0.9 mg kg^-1^, mainly comprising Fe, Si, Al, Li, B, and Cu.

#### 3.4.4. Lipids and carbohydrates

Archaea SCP/PHBV had a low lipid content of 0.93 ± 0.02 g kg^-1^; it mainly consisted of palmitic acid, stearic acid, 1-hexadecanol, and 1-tetracosanol, as well as a low level of carbohydrate content, 3.0 ± 0.2 g kg^-1^; it consisted of arabinose, galactose, and glucose.

#### 3.4.5. Amino acid profile

The amino acid profile of archaea SCP/PHBV (Table 3) consists of 17 amino acids. Tryptophan remained undetectable under the analysis conditions, whereas both asparagine and glutamine were incorporated within the measured values of aspartic acid and glutamic acid, respectively. The archaea SCP/PHBV comprised 358.0 ± 3.9 g kg^-1^ of amino acids, 146.7 ± 4.8 g kg^-1^ of essential amino acids (Hou & Wu, 2018), 211.3 ± 1.0 g kg^-1^ of non-essential amino acids, and 71.2 ± 0.7 g kg^-1^ of branch chain amino acids. According to the Food and Agriculture Organization of the United Nations (FAO) (FAO, 2010), the total amino acids in rolled oats, 101 g kg^-1^, lentils, 16.9 g kg^-1^, wheat, 86 g kg^-1^, peas, 170 g kg^-1^, and kidney beans, 165 g kg^-1^, are lower than those of archaea SCP/PHBV, 358 g kg^-1^, whereas casein, 775 g kg^-1^, has a higher amount of total amino acids. In addition, the essential amino acids in rolled oats, 37 g kg^-1^, lentils, 67 g kg^-1^, wheat, 28 g kg^-1^, peas, 68 g kg^-1^, and kidney beans, 70 g kg^-1^, are lower than those in archaea SCP/PHBV, 146.7 g kg^-1^, whereas casein, 347 g kg^-1^, has a higher amount of essential amino acids.

**Table 3.** Amino acid profile. The amount of amino acid (g kg^-1^, mean ± SD, *n* = 3) in *H. mediterranei* cell dry samples. The results were expressed as means of triplicate ± standard deviation. Tryptophan was not detected.

| No. | Amino acid | Amount (g kg<sup>-1</sup>) |
| --- | --- | --- |
| 1 | Alanine | 23.3 ± 0.2 |
| 2 | Arginine | 24.9 ± 0.3 |
| 3 | Aspartic acid | 45.5 ± 0.2 |
| 4 | Cysteine | 2.9 ± 0.1 |
| 5 | Glutamic acid | 55.7 ± 0.4 |
| 6 | Glycine | 18.1 ± 0.3 |
| 7 | Proline | 13.2 ± 1.2 |
| 8 | Serine | 13.9 ± 0.8 |
| 9 | Tyrosine | 13.7 ± 0.3 |
| 10 | Histidine | 9.2 ± 0.1 |
| 11 | Isoleucine | 16.8 ± 0.2 |
| 12 | Leucine | 26.9 ± 0.1 |
| 13 | Lysine | 15.9 ± 0.7 |
| 14 | Methionine | 7.6 ± 0.2 |
| 15 | Phenylalanine | 19.0 ± 2.2 |
| 16 | Threonine | 23.8 ± 1.3 |
| 17 | Tryptophan | n.d. |
| 18 | Valine | 27.5 ± 0.4 |
| Total non-essential amino acids | | 211.3 ± 1.0 |
| Total essential amino acids | | 146.7 ± 4.8 |
| Total amino acids | | 358.0 ± 3.9 |
| *In vitro* digestibility | | 0.91 ± 0.02 |
| Limiting amino acid |  | L-Lysine |
| Limiting amino acid score | | 0.86 ± 0.01 |
| Protein digestibility-corrected amino acid score (PDCAAS) | | 0.78 ± 0.02 |
| Branch chain amino acids | | 71.2 ± 0.7 |

n.d.: not detected.

In summary, the archaea SCP/PHBV matrix exhibits substantial protein content and quality. It consists of 357.9 g kg^-1^ of proteins and 360 g kg^-1^ of PHBV as the major constituents, whereas ash, 201.5 g kg^-1^, carbohydrates, 3.0 g kg^-1^, lipids, 0.93 g kg^-1^, and moisture, 76.7 g kg^-1^, contributed to SCP/PHBV’s diverse composition (Fig. 3).

**Fig. 3.**
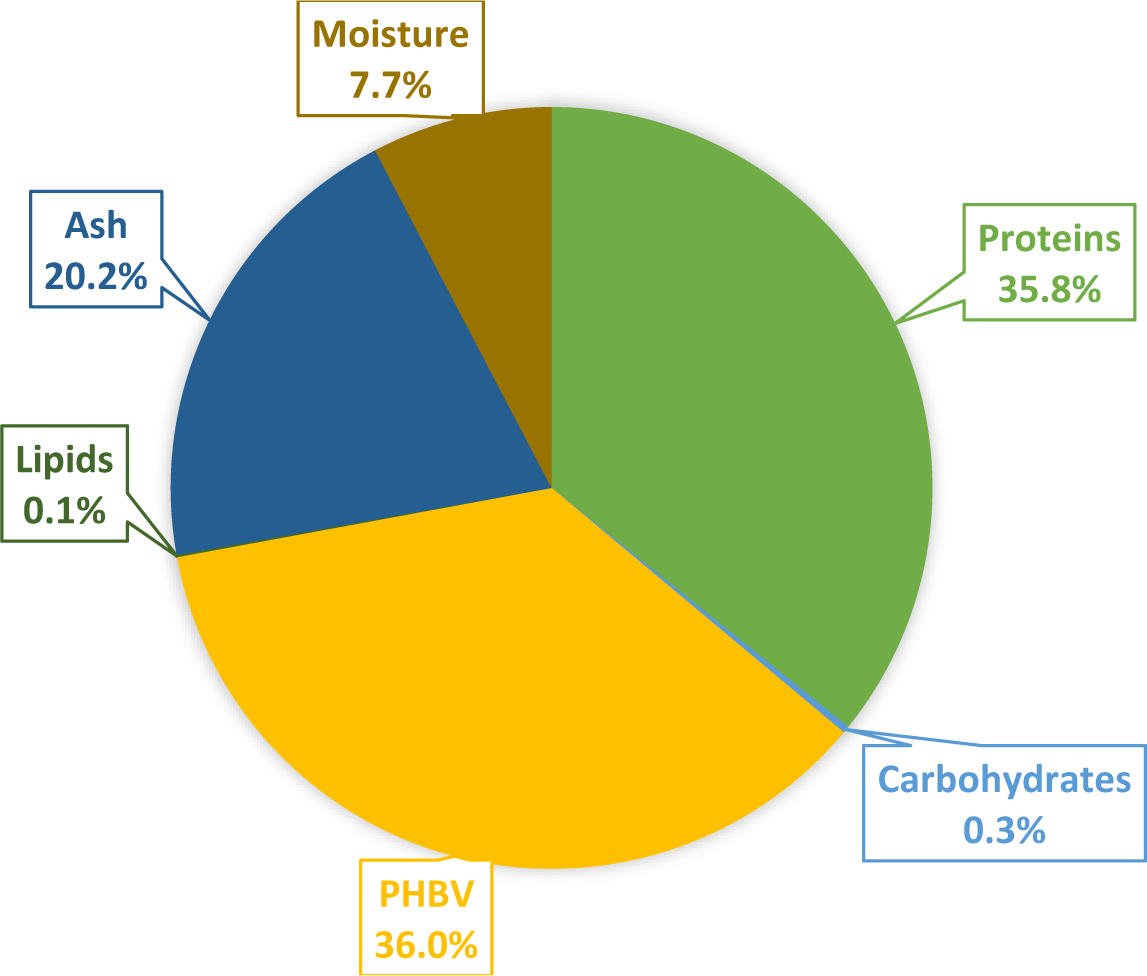
The nutritional profile of SCP/PHBV from *H. mediterranei*: Proteins, PHBV polymer, lipids, carbohydrates, ash, and moisture.

### 3.5. Properties and structural analysis of the archaea SCP/PHBV matrix

#### 3.5.1. Bulk density, true density, and the water- and oil-holding capacities

The peachy-pink color and yeast-like odor of the archaea SCP/PHBV can be attributed to its microbial origin. The SCP/PHBV matrix displayed a bulk density of 0.95 ± 0.01 g cm^-3^, compared to 0.82 g cm^-3^ for yeast biomass and 0.76 g cm^-3^ for wheat flour (Razzaq et al., 2020). In addition, the SCP/PHBV matrix exhibited a true density of 1.58 ± 0.03 g cm^-3^, a water-holding capacity of 2.09 ± 0.01 g/g, and an oil-holding capacity of 0.71 ± 0.06 g/g. These properties indicate that the archaea SCP/PHBV matrix is relatively compact in its solid form and denser when considering its particle arrangement. The water- and oil-holding capacity data are crucial for the future processing of the SCP/PHBV matrix, where moisture and fat retention are essential for stabilizing food formulation.

#### 3.5.2. FTIR analysis of PHBV and SCP/PHBV matrix

FTIR analysis of the SCP/PHBV matrix revealed the characteristic vibration band of the ester carbonyl bond (C=O) at 1724-1734 cm⁻¹ and the stretching band of the C-H bond (CH3) at 2930 cm^-1^, suggesting the presence of PHBV polymer in the SCP/PHBV matrix. The 1628 cm⁻¹ and 1527 cm⁻¹ bands correspond to the protein amide I and II vibrations (Fig. 4). Other peak frequency assignments, such as 1179 cm^-1^ and 978 cm^-1^ for C-O-C vibration, 1054 cm^-1^ for C-O-C and C-C stretching as well as C-O-H bending, were also observed and match the values in the literature (Alsafadi & Al- Mashaqbeh, 2017).

**Fig. 4.**
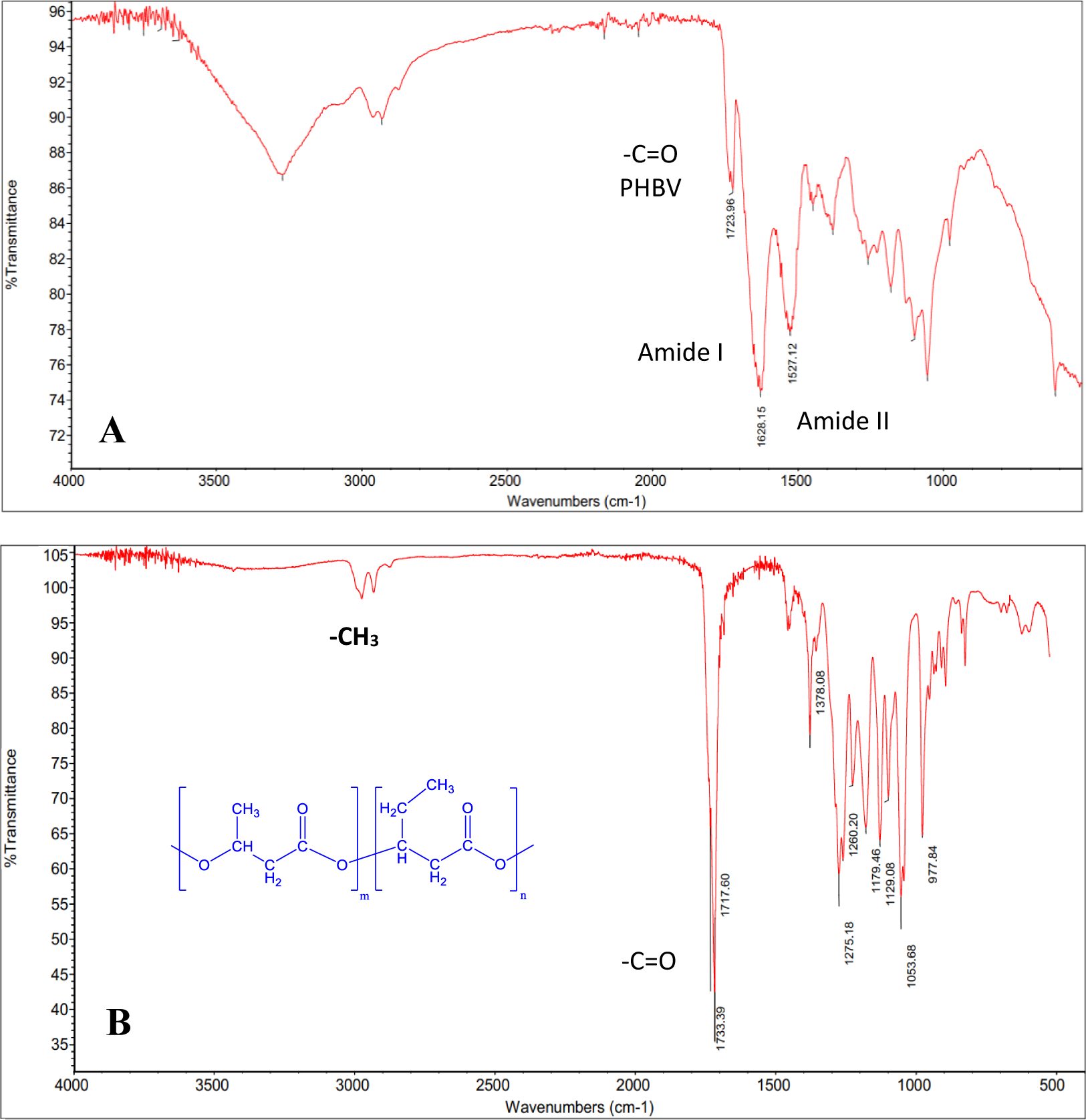
Fourier-Transform Infrared Spectroscopy (FT-IR) spectrum of (A) *H. mediterranei* SCP/PHBV cultivated on BW and (B) pure PHBV extracted with chloroform.

#### 3.5.3. NMR analysis of PHBV

The ^13^C NMR spectra of the pure PHBV polymer (Fig. 5) displayed nine singlet signals as follows: at 169.4 ppm for the two **<u>C</u>**=O carbons, 72.1 ppm for the -**<u>C</u>**H-CH2CH3 carbon, 67.8 ppm for the -**<u>C</u>**H-CH3 carbon, 41.0 ppm and 39.0 ppm for the two-**<u>C</u>**H2-C=O carbons, 27.1 ppm for the -**<u>C</u>**H2-CH3 carbon, 20.0 ppm for the −CH-**<u>C</u>**H3 carbon, and 9.6 ppm for the −CH2-**<u>C</u>**H3 carbon. The ^1^H NMR spectra of the PHBV polymer exhibited seven signals as follows: a sextet and quintet at 5.21-5.26 ppm for the −C**<u>H</u>**- hydrogens, two doublets of doublet at 2.42-2.61 ppm for the −C**<u>H</u>2**-CO- hydrogens, a quintet at 1.58 ppm for the −C**<u>H</u>2**-CH3 hydrogens, a doublet at 1.24-1.26 ppm for the −CH-C**<u>H</u>**3, and a triplet at 0.83 ppm for the −CH2-C**<u>H</u>**3 hydrogens (Ghosh et al., 2019). The ^1^H- and ^13^C-NMR spectra of the extracted PHBV polymer (Fig. 5) confirmed the structure of a copolymer of 3-hydroxybutyrate and 3-hydroxyvalerate with a composition of 91:9 mol%, as determined by GC-MS.

**Fig. 5.**
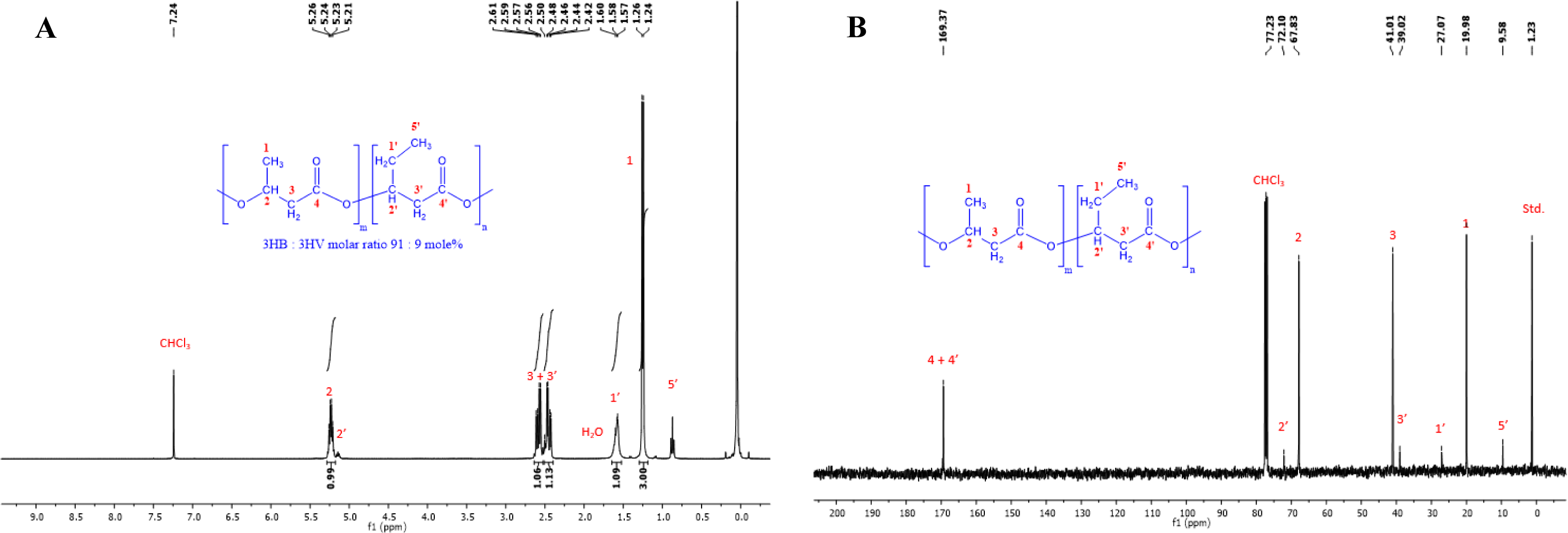
(A) Proton nuclear magnetic resonance spectroscopy (^1^H NMR) (400 MHz, CDCl3) spectra of PHBV extracted by chloroform from the cell dry mass of *H. mediterranei* cultivated in bread waste (BW). (B) Carbon nuclear magnetic resonance spectroscopy (^13^C NMR) (400 MHz, CDCl3) spectra of PHBV extracted by chloroform from the cell dry mass of *H. mediterranei* cultivated in bread waste (BW).

#### 3.5.4. GPC analysis of PHBV

The GPC analysis conducted on the extracted PHBV polymer from *H. mediterranei* SCP/PHBV (Fig. 6) revealed that the *Mw* = 3827 kDa and *Mn* = 3140 kDa values of PHBV produced from BW were notably higher than those of PHBV produced from glucose (*Mw* = 2564 kDa, *Mn* = 2115 kDa) but had a similar polydispersity index (*PDI = Mw/Mn*) of 1.212 and 1.219, respectively, which suggests the production of homogeneous PHBV polymers in both cases. The high *Mw* value of PHBV reflects its improved mechanical properties, including increased tensile strength and durability. Sato et al. reported that the *Mw*, *Mn*, and *PDI* values of PHBV produced by *H. mediterranei* in different media are 1722-5280 kDa, 800-3500 kDa, and 1.6-2.2, respectively (Sato et al., 2021). This study demonstrated that under low concentrations of amino acid sources, 0.1-1 g L^-1^, the *Mw* = 5500 kDA and *Mn* = 3500 kDa values of the produced PHBV were relatively high, possibly due to the amount of PHBV synthase produced under such conditions.

**Fig. 6.**
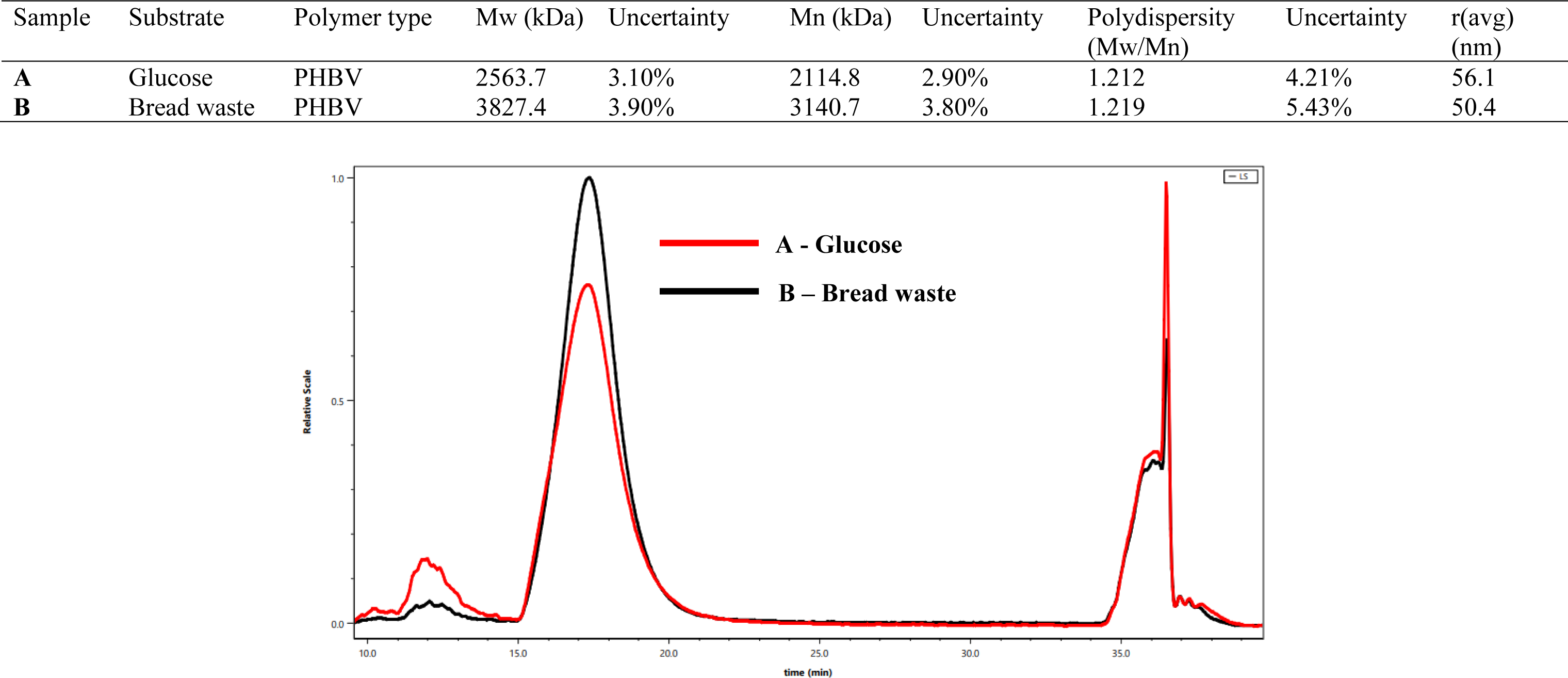
Gel permeation chromatography (GPC) of PHBV extracted by chloroform from the cell dry mass of *H. mediterranei* cultivated glucose (red). (B) Gel permeation chromatography (GPC) of PHBV extracted by chloroform from the cell dry mass of *H. mediterranei* cultivated in bread waste (BW) (black).

#### 3.5.5. TGA and DSC analysis

The thermal stabilities of SCP/PHBV, isolated PHBV, and isolated protein were investigated by TGA and DSC (Fig. 7). PHBV exhibited thermal stability with an initial weight decrease at 258 °C, *Td* at 277 °C, and *Tm* at 274 °C. The protein underwent thermal degradation starting at 272 °C, with a significant structural change at 312 °C, *Td* at 349 °C, and a cross-linking transition at 370 °C. SCP/PHBV experienced thermal degradation at 244 °C, featuring a notable structural alteration at 255 °C, and it reached a substantial decomposition at 269 °C, along with a *Tm* value of 258 °C and a cross-linking point at 538 °C. Using TGA, Singha et al. examined the interaction between microbial SCP and bioplastic films (Singha et al., 2021). Their findings indicated a shift towards higher temperatures, suggesting that the thermal stability behavior was affected by the development of intermolecular interactions between glycerol and the amino acid groups, especially when subjected to increased temperatures and extended pressing durations.

**Fig. 7.**
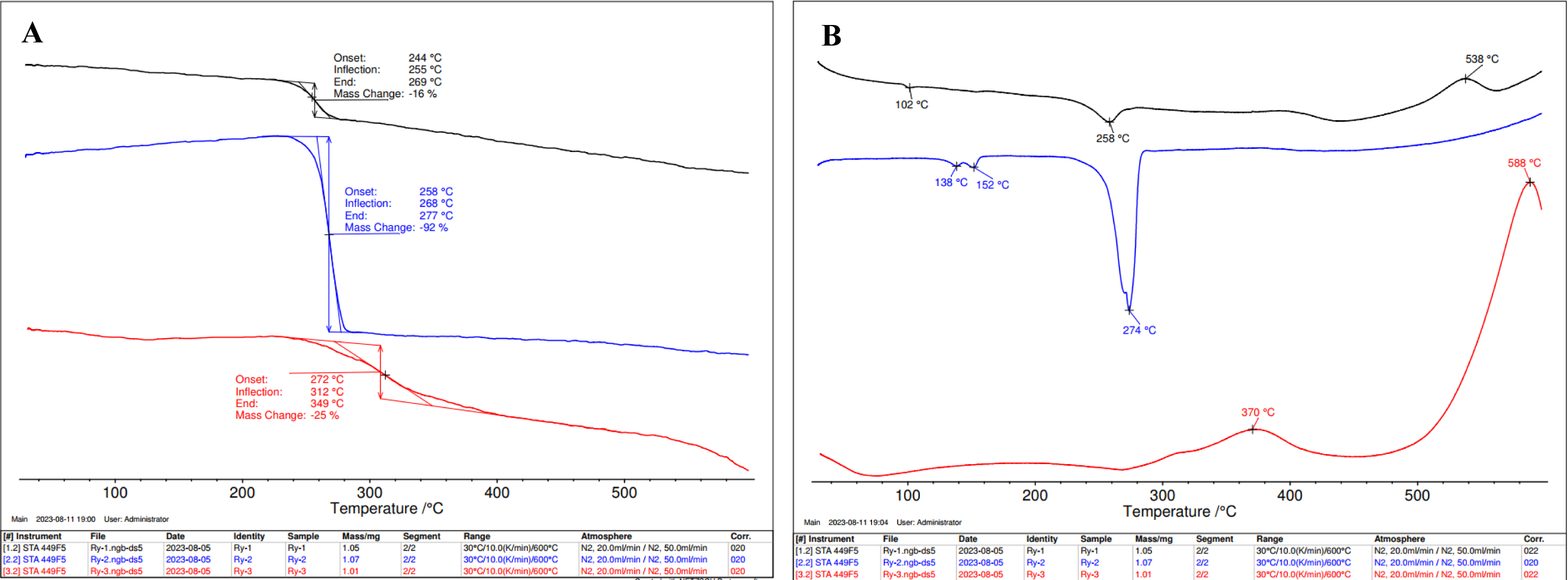
(A) Thermo gravimetric analysis (TGA) and (B) Differential scanning calorimetry (DSC) of *H. mediterranei* SCP/PHBV (black), PHBV (blue), and protein (red) extracted from *H. mediterranei* cultivated on bread waste (BW).

### 3.6. Protein quality of archaea SCP/PHBV

SCP’s protein quality was assessed by evaluating its amino acid composition and its *in-vitro* protein digestibility-corrected amino acid score (PDCAAS). The calculated parameters for *H. Mediterranei* SCP/PHBV were as follows: the *in-vitro* digestibility was 0.91, the first limiting amino acid (L-lysine) score was 0.86, the essential amino acid ratios ranged from 0.86 to 2.45, the PDCAAS was 0.78, and the crude protein percentage values were 35.8% (Table 3). According to the FAO recommendation (FAO, 2010), the PDCAAS values of high- and excellent-quality protein should be higher than 0.75 and 1.00, respectively (Zeng, Chen, Zhang, & Wang, 2022). Animal protein sources such as casein have PDCAAS values equal to 1 (Qin, Wang, & Luo, 2022). Plant protein has lower PDCAAS values that range from 0.39 to 1.00, for instance, and the soy protein PDCAAS value is 0.92 (Rutherfurd, Fanning, Miller, & Moughan, 2015), whereas the almond PDCAAS value is 0.39 (Weindl et al., 2020). Edible mycoproteins have a PDCAAS value of 0.35-0.70 (Ahmad, Farooq, Alhamoud, Li, C & Zhang, 2022), and yeast protein concentrate has a range of 0.82 to 0.90 (Ariëns et al., 2021). Regarding algae, the PDCAAS values range from 0.08 to 0.69 (Zeng et al., 2022). Therefore, archaea SCP/PHBV derived from *H. Mediterranei* has a relatively high protein quality, as indicated by its PDCAAS value (0.78).

### 3.7. Valorization of bread waste

The current study aimed to demonstrate the microbial bioconversion of BW hydrolysate to the SCP/PHBV matrix by *H. mediterranei*. The main strategy regarding a substrate used to produce microbial biomass containing high-quality protein is to consider low-grade waste material (Carranza-Méndez et al., 2022). BW is regarded as a potential carbon/nitrogen source. The global annual bread production is >100 million tonnes (Narisetty et al., 2021). Owing to a short shelf-life and the overproduction of bread, approximately 10% (∼ 10 million tonnes) of the bread produced globally is discarded; it amounts to around 24 million slices of bread every day, representing 27- 31% of the total food waste mass (Montemurro et al., 2022). In the UK, the largest bread consumer in Europe, approximately 0.3 million tonnes of bread is wasted annually (Jung et al., 2022). In Israel and the USA, within the grain and legume category, the waste rate stands at approximately 14% (0.17 million tonnes) and 25% (0.31 million tonnes), respectively (Philip, Hod-Ovadia, & Troen, 2017). Disposal of BW without its valorization could result in the loss of resources (Jung et al., 2022). Hydrogen, ethanol, lactic acid, succinic acid, lipids, and PHA are examples of high-value products generated by the microbial fermentation of BW (Montemurro et al., 2022). Therefore, BW can potentially be utilized as a valuable and sustainable carbon/nitrogen source that would contribute to a sustainable coproduction of archaea SCP/PHBV while simultaneously addressing waste reduction and resource optimization challenges.

The carbon footprint of 1000 g of BW was estimated to be 1555 g of CO2 and 350 mL of CH4 emissions (Carlsson &Uldal, 2009). Using BW (10 million tonnes worldwide) in microbial fermentation by *H. Mediterranei* to produce SCP/PHBV, rather than allowing it to decompose in landfills, could potentially reduce CH4 emissions by 3.5 million tonnes/year while generating 6.5 million tonnes/year of CO2. Furthermore, besides the financial and bioresource losses, BW, like any other food waste, causes significant damage to the broader environment by contributing to global warming, acidification, and eutrophication (Ben Rejeb, Charfi, Baraketi, Hached, & Gargouri, 2022). Therefore, the reduction, recycling, or valorization of food waste can potentially save billions of dollars in global economic value, safeguard invaluable bioresources, and prevent the release of millions of tons of greenhouse gases into the atmosphere.

### 3.8. Applications and functional benefits of SCP/PHBV matrices

The versatility of SCP/PHBV matrices is evident in its wide array of applications, spanning from animal feed and human food to industrial processes. Opting to directly utilize SCP/PHBV matrices offers several advantages, considering factors such as cost, nutritional requirements, sustainability, and product attributes. SCP/PHBV matrix, with its whole-cell composition, typically preserves a richer array of nutrients and bioactive compounds compared with extracted protein, rendering it highly beneficial for both human and animal consumption. Furthermore, the holistic nature of SCP/PHBV matrices has the potential for added functional benefits, such as fiber content and unique cell components, which can prove advantageous in specific food products and applications. Direct utilization of SCP/PHBV streamlines processes, consequently reducing the reliance on expensive extraction and purification methods, ultimately establishing it as a cost-effective protein source. Since it is environmentally conscientious, this approach often entails fewer processing steps and lower energy consumption than do protein extraction processes, thus contributing to sustainability goals. Furthermore, the SCP/PHBV matrix retains the natural flavors and textures of the organism, a desirable quality in food products where taste and mouthfeel are paramount. However, archaea SCP/PHBV matrices must undergo comprehensive safety testing, including toxicity assessments and allergenicity testing, to ensure their suitability and safety for human consumption.

## 4. Conclusions

This study investigated the SCP/PHBV matrix production by *H. mediterranei* using BW as a substrate for the first time. Importantly, this research revealed that BW samples varied significantly in their nutritional composition, with significant differences in carbohydrates, lipids, proteins, dry weight, and ash content. Despite this variability, all BW samples were rich in nitrogen, making them suitable for microbial protein production. Moreover, this study identified critical factors affecting *H. mediterranei* growth, such as pH, red sea salt concentration, and BW concentration. Under optimized conditions, a mixture of 30 g L^-1^ BW hydrolysate and 200 g L^-1^ of red sea salt produced a maximum SCP/PHBV concentration of 8.0 ± 0.1 g L^-1^. The SCP/PHBV matrix exhibited unique properties, including high protein content and remarkable mineral content due to its high salinity origin. The SCP/PHBV matrix displayed good protein quality, as indicated by its amino acid composition and PDCAAS, making it a promising protein source for various applications. Utilizing BW as a substrate for SCP/PHBV production could significantly reduce CH4 emissions and CO2 generation from BW, thus contributing to waste reduction and resource optimization while addressing environmental challenges. Our long-term goal is to develop a new strategy to fabricate an SCP/PHA matrix suitable for high moisture extrusion of meat analogs. Although this study provides promising insights into the production of SCP/PHBV matrices by *H. mediterranei*, further research and safety testing are required to fully assess its suitability for human consumption and agriculture purposes.

## CRediT authorship contribution statement

**Razan Unis**: Investigation, Methodology, Formal analysis, Writing - original draft, and Review & editing. **Rima Gnaim**: Investigation, Data curation, and Formal analysis. **Mrinal Kashyap**: Investigation, Data curation, and Formal analysis. **Olga Shamis**: Formal analysis. **Nabeel Gnayem**: Formal analysis. **Michael Gozin**: Supervision. **Alexander Liberzon**: Supervision. **Jallal Gnaim**: Supervision, Writing - review & editing. **Alexander Golberg**: Supervision, Conceptualization, Review.

## Declaration of Competing Interest

The authors declare that they have no known competing financial interests or personal relationships that could have appeared to influence the work reported in this paper.

## Acknowledgments

J.G., A.G., and A.L. thank the Ministry of Science & Technology, the Ministry of Agriculture, and the Good Food Institute (GFI).

## Statement of informed consent

No conflicts, informed consent, human or animal rights are applicable.

## Declaration of authors’ agreement to authorship and submission

All authors have agreed to the authorship and the submission of this manuscript.

## Appendix A. Supplementary data

Word document.

## Notes

### Competing Interest Statement

The authors have declared no competing interest.

