## Supplementary for "Production of single-cell-protein (SCP) / poly(3-hydroxybutyrate-co-3-hydroxyvalerate) (PHBV) matrices through fermentation of archaea *Haloferax mediterranei*"

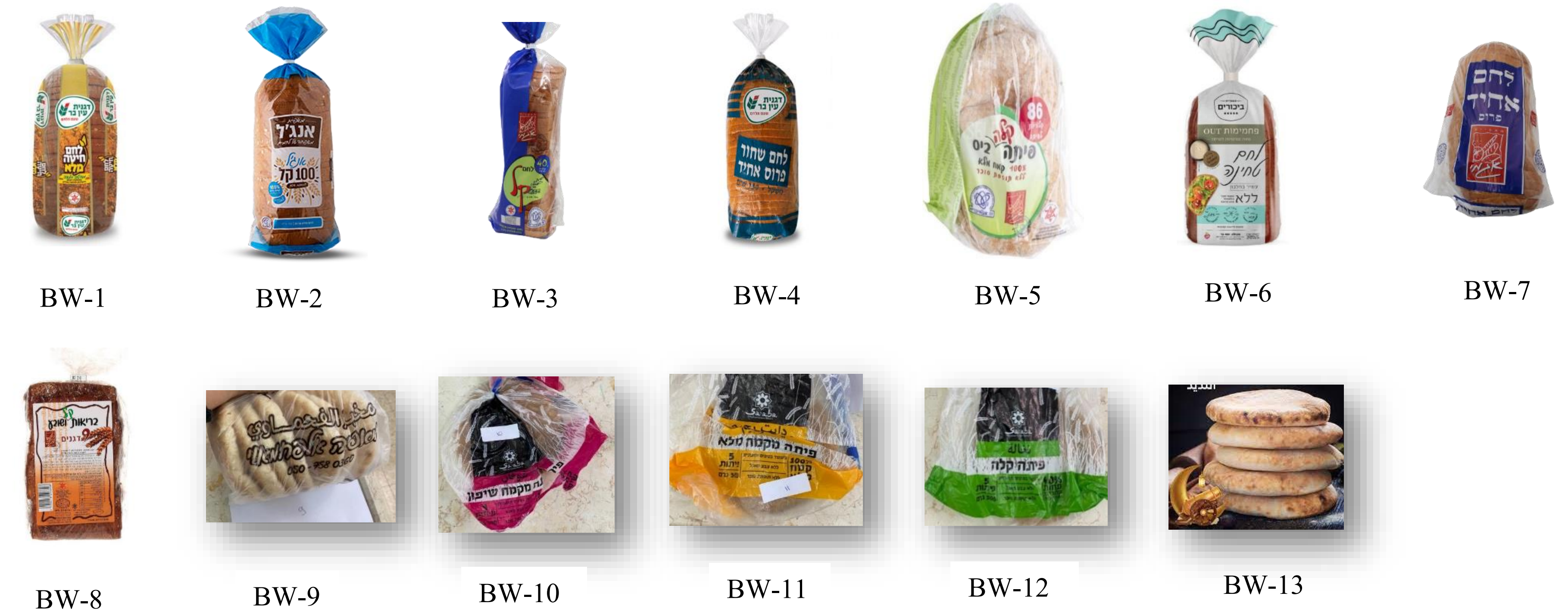


Supplementary Figure 1. Bread samples: BW-1 Whole wheat bread, Deganit Ein Bar, Israel; BW-2 Sliced whole meal bread, Angel - Israel; BW-3 Light bread, Agami, Israel; BW-4 Sliced black bread, Deganit Ein Bar, Israel; BW-5 Light bite pita, Agami, Israel; BW-6 Flax bread - first fruits, Israel; BW-7 Sliced uniform white bread, Israel; BW-8 Cereal bread, Agami, Israel; BW-9 White pita, Alfachamaoui bakery, Israel; BW-10 Rye flour pita, Saaba, Israel; BW-11 Wholemeal pita bread, Saaba, Israel; BW-12 Light pita, Saaba, Israel; BW-13 Light pita, Saaba, Israel.
